# A class of COPI adaptors regulates processing of transmembrane receptors by reinforcing their Golgi retention

**DOI:** 10.1101/2025.04.11.648407

**Authors:** Jinghu Gao, Chengge Cong, Yun Xiang, Yanfang Wu, Bing Yan, Qinyu Jia, Zhong-ping Yao, Yusong Guo, Junjie Hu

**Affiliations:** Department of Genetics and Cell Biology, College of Life Sciences, Nankai University, Tianjin 300071, China; Key Laboratory of Biomacromolecules, Institute of Biophysics, Chinese Academy of Sciences, Beijing 100101, China; Division of Life Science, Hong Kong University of Science and Technology, Hong Kong, China; University of Chinese Academy of Sciences, Beijing 100101, China; State Key Laboratory of Chemical Biology and Drug Discovery, Research Institute for Future Food, Research Centre for Chinese Medicine Innovation, and Department of Applied Biology and Chemical Technology, The Hong Kong Polytechnic University, Hong Kong, China

**Author notes:** These authors contributed equally (Jinghu Gao, Chengge Cong). To whom correspondence should be addressed (J.H.), (Y.G.), and (J.G.).

## Abstract

Many transmembrane (TM) receptors undergo essential post-translational modification in the Golgi prior to their delivery to the plasma membrane. Whether and how the passage and accompanied modification of these receptors across the Golgi is controlled remains unclear. Here, we show that leptin receptor overlapping transcript (LEPROT) and LEPROT-like 1 (LEPROTL1) regulate receptor activation by securing their sufficient Golgi retention. LEPROTs localize to cis and medial Golgi in a COPI-dependent manner. Deletion of LEPROTs in cells causes expedited release of receptors transiting through the Golgi, and leakage of some Golgi-enriched proteins into the endomembrane compartments. LEPROTs interact directly with COPI coats and simultaneously engage a variety of integral membrane proteins with relatively long TM domains at acidic pH. Loss of LEPROTs dysregulates receptor activity, including that of EGFR and TFRC, due to defective modification. Collectively, LEPROTs serve as a class of COPI adaptors for TM receptors, ensuring adequate preparation, which is vital for subsequent action on the plasma membrane.

## INTRODUCTION

In eukaryotic cells, the Golgi apparatus is a stack of cisternal structures that mediate protein processing and sorting (1–3). During interphase, Golgi are localized to the perinuclear region in proximity to the microtubule organizing center (MTOC) (4). From cis to trans, each layer of Golgi contains designated markers, mostly rod-like Golgins that help maintain the morphology and function of Golgi, and signature enzymes that are responsible for regulated proteolysis or further modification of proteins initially synthesized in the endoplasmic reticulum (ER) (5, 6). After reaching the trans Golgi network (TGN), proteins are considered fully mature and are sorted and packed into different vesicles, which are subsequently delivered to various compartments of the endomembrane system, such as lysosomes, endosomes, and the plasma membrane (PM) (7, 8).

The protein trafficking in and out of Golgi is mediated by a set of coated vesicles. The COPII vesicles deliver cargo from the ER to Golgi, the COPI vesicles convey retrograde transport from Golgi back to the ER and transport within Golgi, and clathrin-coated vesicles (CCVs) export proteins from the TGN (2, 9, 10).

Compromised COPII or clathrin assembly causes defects in exporting cargo proteins out of the ER or the Golgi, respectively, whereas defective COPI trafficking expedites protein release from the Golgi (11–13), specifically within the framework of the cisternal progression model (14).

The vesicle formation process usually involves initiating complex, coats, cargo adaptors/receptors and various cargo clients. Small GTPases, including Sar1 and Arf1, initiate vesicle formation (15, 16). The coats (i.e., Sec23/24 and Sec13/31 for COPII, coatmer complex for COPI, and clathrin for CCVs) not only stabilize the curvature needed in vesicle budding, but also engage cargo for their precise sorting (17–19).

Cargo adaptors interact simultaneously with cargo and the corresponding coats (11). Enrichment of cargo proteins into CCVs is facilitated by adapter protein (AP) complexes (8) and the Golgi-localized, γ-ear-containing, Arf-binding proteins (GGAs) (20). The COPII subunit, Sec24, functions as a classic cargo receptor that mediate the packaging of cargo proteins into ER-derived vesicles (21). Similarly, COPI clients need to interact directly with the coat or indirectly via cargo adaptors (22). Vps74p, which has been identified as a COPI adaptor in yeast cells, maintains localization of Golgi glycosyltransferases by packaging them into COPI-coated vesicles (23, 24). The mammalian homolog of Vps74p, GOLPH3 also interacts with COPI coats and the cytosolic tails of some Golgi glycosylation enzymes and mediates their Golgi retention (13, 25, 26).

Transmembrane (TM) receptors undergo various processing or modification steps in Golgi before heading towards the PM. The most well-known modification is glycosylation. Golgi-resident glycosyltransferases edit oligosaccharides that are initially added in the ER or add them freshly as proteins pass through the Golgi. Golgi-based glycosylation includes O-linked glycosylation and further modification of N-glycans, with sugars, such as mannose, galactose, or glucose, added one at a time (5). In addition, incorporated sugars or protein residues can be sulfated in Golgi. Finally, protein acylation, similar to palmitoylation, occurs frequently in Golgi (27, 28). In combination, these modifications are critical to the activity of TM receptors. As mentioned above, the known COPI adaptors only serve Golgi-resident enzymes. It remains elusive whether and how the passage of TM receptors through the Golgi is controlled by specific cellular factors. Previous studies for regulating the activities of TM receptors emphasize the retrograde route, which includes endocytosis and Golgi retrieval via recycling pathways. TM receptors can be removed from their post via these pathways. A class of integral membrane proteins called endospanins, also known as leptin receptor overlapping transcript (LEPROT) and LEPROT-like 1 (LEPROTL1), regulate the recycling pathway of leptin receptor after its endocytosis (29). Whether TM receptors can be regulated during the antegrade maturation pathway, and if so, whether proteins like LEPROTs are involved, is not clear. Here, we demonstrate that LEPROTs interact with COPI coats, delay the Golgi release of TM receptors and, in doing so, regulate the modification of these receptors and subsequently affect their activity.

## RESULTS

### LEPROTs localize to Golgi and function in Golgi

In an effort to identify reticulon-homology domain (RHD)-containing proteins, which possess two consecutive transmembrane hairpins (TMHs) and potentially play a role in membrane curvature stabilization (30), we found LEPROT and LEPROTL1. To confirm that they are indeed integral membrane proteins, we performed membrane fractionation. Endogenous LEPROT or LEPROTL1 with knocked-in C-terminal 2xHA behaved as an integral membrane protein. Both proteins were found in the membrane fractions and were resistant to alkaline extraction (**Fig. 1a**). The LEPROTs are predicted to have four TM domains, with very short loops connecting TM1-TM2 and TM3-TM4, hence two TMHs (**Fig. 1b**). The membrane topology of LEPROTs has been tested previously (29), and was confirmed here with a different method (**Fig. S1a,b**).

**Fig. 1.**
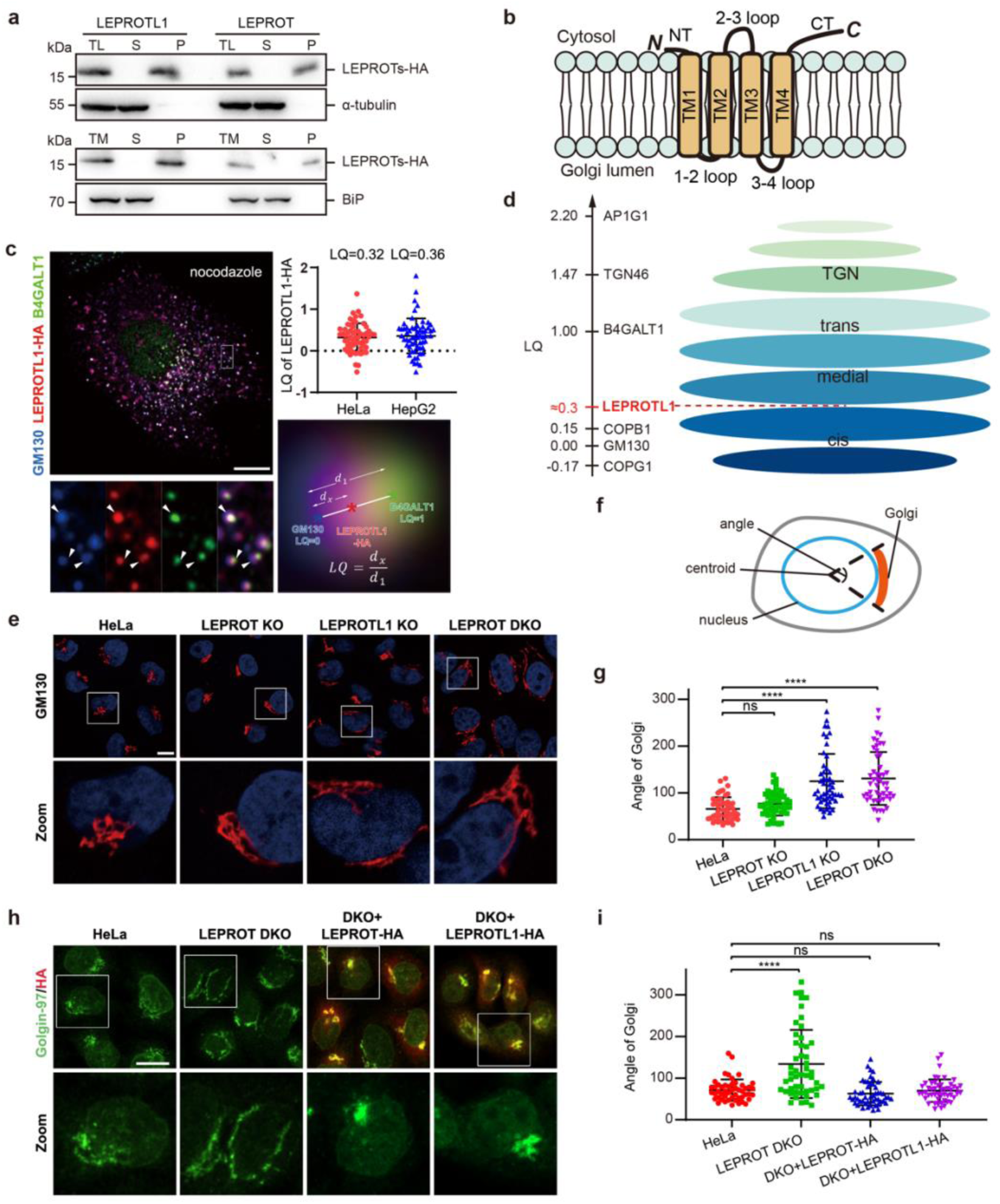
LEPROTs control Golgi morphology. (**a**) Total lysate (TL) of LEPROTs-HA knockin HeLa cells was separated into soluble (S) and pelleted membrane (P) fractions and then immunoblotted for HA or the cytosolic protein α-tubulin. The total membrane (TM) fractions were alkaline-extracted and S and P fractions immunoblotted for HA and the soluble lumenal protein BiP. (**b**) Topological diagram of LEPROTL1. Given the sequence conservation between LEPROT and LEPROTL1, the same diagram would be applicable to LEPROT. (**c**) HepG2 cells transiently expressing LEPROTL1-HA (red) were treated with nocodazole and processed for immunofluorescence labeling of endogenous B4GALT1 (green) and GM130 (blue). Arrowheads in the zoomed region indicate overlap of GM130, LEPROTL1-HA, and B4GALT1. The bottom right Golgi mini-stack schematically illustrates centers of mass (denoted by asterisk, *) and calculation of the LQ. The LQ scatter plots for HeLa and HepG2 cells are at the top right. The values shown in the scatter plots are mean ± SD (n=60). Scale bar = 10 μm. (**d**) A schematic diagram of an average Golgi mini-stack. LQs of various Golgi proteins, including LEPROTL1, are overlaid onto the y-axis. (**e**) Confocal images of the Golgi marked by GM130 in wild-type, LEPROT KO, LEPROTL1 KO, and LEPROT DKO HeLa cells. Nuclei were indicated using DAPI (blue). Scale bar = 10 µm. (**f**) Diagram of the measurement standards for the angle of Golgi. (**g**) Quantification of the angle of Golgi in (e) based on confocal images. Values are mean ± SD; n=50, each dot represents one cell. ****p<0.0001, two-tailed Student’s t-test. (**h**) LEPROT DKO HeLa cells were generated by lentiviral infection to stably express LEPROT-HA or LEPROTL1-HA. Cells were fixed and stained using Golgin-97 and HA antibodies. (**i**) Quantification of the angle of Golgi in (h) based on confocal images. Values are mean ± SD; n=50, each dot represents one cell. ****p<0.0001, ns: not significant, two-tailed Student’s t-test. Scale bar = 10 μm.

Next, we tested the intracellular localization of LEPROTs. Previous studies have reported that LEPROTL1 (also known as endospanin-2) is mostly enriched in the Golgi, but LEPROT (also known as endospanin-1) is often found in endosomes with some Golgi localization (29). Consistently, we found that LEPROTL1-HA was predominantly enriched in the Golgi with additional puncta overlapping with endosomal marker EEA1, but not lysosomal marker LAMP1 (**Fig. S1c**). However, the same results were obtained when LEPROT-HA was analyzed (**Fig. S1d**), suggesting uncertainty in the endosomal enrichment of LEPROT. We then detected localization of endogenous LEPROTL1 in various cells using anti-LEPROTL1 antibodies for immuno-staining and confirmed the predominant Golgi residency (**Fig. S1e**).

To determine the precise positioning of LEPROTs within the Golgi apparatus, we performed the previously established <u>G</u>olgi protein <u>L</u>ocalization by Imaging centers of <u>M</u>ass (GLIM) (32–34). Briefly, Golgi were dissembled into mini stacks by nocodazole treatment, and the localization of LEPROTL1-HA was referenced to cis-Golgi marker GM130 and trans-Golgi marker B4GALT1 (**Fig. 1c**). The calculated localization quotient (LQ) indicated that LEPROTL1 resides between the cis and medial Golgi (**Fig. 1d**). These results suggest that LEPROTs most likely function in Golgi.

To test whether LEPROTs influence Golgi structures, we generated CRISPR/Cas9-mediated knockout (KO) lines in various cells. In the process of verifying these cells at the protein level, we found that endogenous LEPROTL1 could barely be detected using conventional immuno-blotting preparation even though the antibody worked efficiently with overexpressed protein. Only when cells were first permeabilized with digitonin, lysed under low pH, and denatured without heating was LEPROTL1 readily seen (**Fig. S1f**). These observations indicate that LEPROTs are aggregation-prone. It also explains why these factors are rarely detected in previous proteomics studies of the endomembrane system. Using the optimized method, we confirmed the successful deletion of LEPROTL1 in a series of KO cell lines (**Fig. S1g**).

Golgi in HeLa cells exhibited a typical peri-nuclear cluster and LEPROT KO caused marginal changes (**Fig. 1e**). In contrast, both the LEPROTL1 KO and double KO (DKO) cells displayed a Golgi ribbon extension around the nucleus (**Fig. 1e**), which can be scored by measuring the perinuclear angle of Golgi (**Fig. 1f,g**). Similar results were obtained when LEPROTs were deleted in COS-7 cells (**Fig. S1h,i**). We then tested whether LEPROT acts in the same manner as LEPROTL1 in the maintenance of Golgi structures. When LEPROT-HA or LEPROTL1-HA was individually reintroduced into DKO HeLa cells, the Golgi ribbon extension was reversed to the same extent as that of the wild-type (**Fig. 1h,i**). These results suggest that LEPROT possibly has similar function or activity as LEPROTL1. Notably, in the single-copy KI of LEPROT or LEPROTL1 in HeLa cells, LEPROTL1 was present at only slightly higher levels than LEPROT (**Fig. 1a**). Therefore, the difference in Golgi phenotype seen with LEPROT or LEPROTL1 KO HeLa cells could not be simply explained by their abundance, implying some functional diversity (see below). Collectively, these results confirm that LEPROTs function in Golgi.

### LEPROTs work with COPI

Given their previously reported activity and our localization analysis, we reasoned that LEPROTs may participate in Golgi-related trafficking and thus tested whether LEPROTs associate with specific coats. When LEPROTL1-HA was immunoprecipitated, endogenous COPI coats, including COPB1 and G1, but not COPII (SEC24C), were readily detected in the precipitates (**Fig. 2a**). To test direct interactions, we synthesized biotinylated peptides derived from the 2-3 loop and C-terminus (CT) and performed pull-down assays. Various endogenous COPI coats were found in the C-terminus, but not the 2-3 loop-related, precipitates (**Fig. 2b**). We also purified one of the COPI components, COPD (**Fig. S2a**), and detected its binding to the CT peptide (**Fig. 2c,d**), but not the 2-3 loop peptide (**Fig. S2b,c**). We noticed that the CT of LEPROTs contained an WxxW motif (**Fig. 2e**), which is known to engage COPD (35). Mutations based on sequence alignment of the C-termini were introduced into the CT peptide and their affinities for COPD compared. Conserved residues proceeding the WxxW motif had a marginal impact on the association, whereas the substitution of either W with A drastically interfered with the binding (**Fig. 2c,d**). These results indicate that LEPROTs directly interact with COPI coats, which fulfill the criteria for serving as a cargo adaptor.

**Fig. 2.**
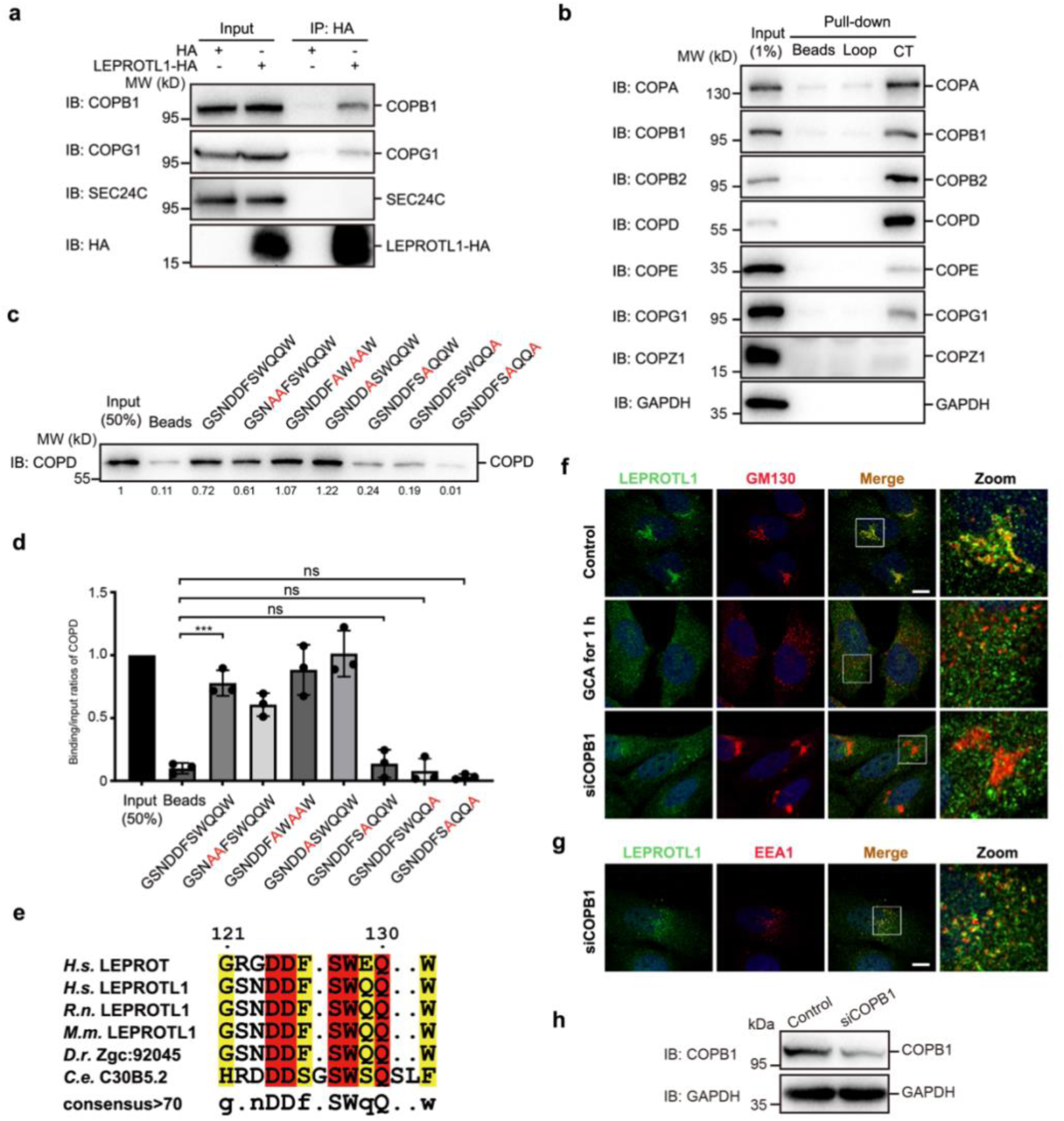
LEPROTs interact with COPI coats. (**a**) Co-immunoprecipitation experiments showing the interaction between LEPROTL1 and COPI complex in HeLa cells. COPII coat SEC24C was used as a negative control. (**b**) Streptavidin beads loaded with biotinylated 2-3 loop or C-terminal LEPROTL1 peptides were incubated with HeLa cell lysate in vitro. Eluted proteins were separated by SDS-PAGE and analyzed by Western blotting using GAPDH and various COPI subunit antibodies. (**c**) Streptavidin beads loaded with biotinylated LEPROTL1 C-terminal or various mutant peptides were incubated with purified COPD in vitro. Eluted proteins were separated by SDS-PAGE and analyzed by Western blotting using COPD antibody. Quantifications of COPD levels are also shown. (**d**) Quantitative data for (d) as mean ± SD (n=3). ***p<0.001, ns: not significant. (**e**) Multispecies alignment of LEPROT C-terminal peptide between *Homo sapiens (H.s.), Rattus norvegicus (R.n.), Mus musculus (M.m.), Danio rerio (D.r.),* and *Caenorhabditis elegans (C.e.)*. (**f,g**) U-2 OS cells treated with negative control or 10 μM Golgicide A for 1 h or transfected with siRNA targeting COPB1 were stained with GM130 or EEA1 antibodies (red) and LEPROTL1 antibodies (green). Nuclei were indicated using DAPI (blue). Scale bar = 10 μm. **(h)** Western blot images showing the knockdown efficiency of COPB1 in (**f,g**).

We then tested whether COPI interferes with the localization of LEPROTs. In U-2 OS cells, in which endogenous levels of LEPROTL1 are relatively high and readily detected by anti-LEPROTL1 staining, LEPROTL1 scattered as puncta throughout the cytosol when cells were treated with Golgicide A (GCA), an inhibitor of Arf GEF GBF1 that blocks COPI activity (36). In these cells, the Golgi was fragmentated, but GM130 puncta marginally overlapped with those of LEPROTL1 (**Fig. 2f**), indicative of the Golgi departure of LEPROTL1. Furthermore, mild depletion of COPB1 (**Fig. 2h**), a component of the COPI coat, resulted in a scattered distribution of LEPROTL1 that did not overlap with GM130 (**Fig. 2f**). Notably, some dispersed LEPROTL1 signal co-localized with endosomal marker EEA1 (**Fig. 2g**), suggesting that some escaped LEPROTs were transported into endosomes in COPI-depleted cells. These analyses indicate that the COPI coat (Fig. 1d, LQ=0.15) is crucial for effectively retaining LEPROTs at the Golgi (Fig. 1d, LQ=0.3), with minimal chances for these proteins escaping this retention and participating in clathrin-mediated trafficking at the TGN (Fig. 1d, AP1G1, LQ=2.2) under physiological conditions. In other words, LEPROTL1 works mainly with COPI in membrane trafficking.

Next, we sought to investigate whether COPI vesicles contain LEPROTs and whether LEPROTs-enriched vesicles carry specific clients. For this purpose, we performed the vesicle formation assay utilizing HEK293T cells transfected with plasmids encoding these proteins linked to a C-terminal HA tag by a poly-glycine linker (LEPROT-HA or LEPROTL1-HA). As expected, LEPROT or LEPROTL1 were readily found in the vesicle fraction when the assay was performed in the presence of cytosol (**Fig. 3a** and **Fig. S3a**). The GTP hydrolysis-defective mutant, SAR1A(H79G), greatly decreased COPII formation, as shown by the diminished abundance of the known COPII export cargo ERGIC53 in the vesicle fraction but had little effect on the appearance of LEPROTs (**Fig. 3a** and **Fig. S3a**). We also tested whether deleting COPI from the cytosol would impair the budding behavior of LEPROTL1. When the CT peptide of LEPROTs was incubated with cytosol extracts prepared from HEK293T cells, the COPI coats were efficiently depleted (**Fig. S3b**). Consistent with the observation in intact cells, more LEPROTL1 was detected in the vesicle fraction (**Fig. S3c**), suggesting failure of its Golgi retention. These findings confirm that COPI coat is important for the retention of LEPROTL1 in the Golgi.

**Fig. 3.**
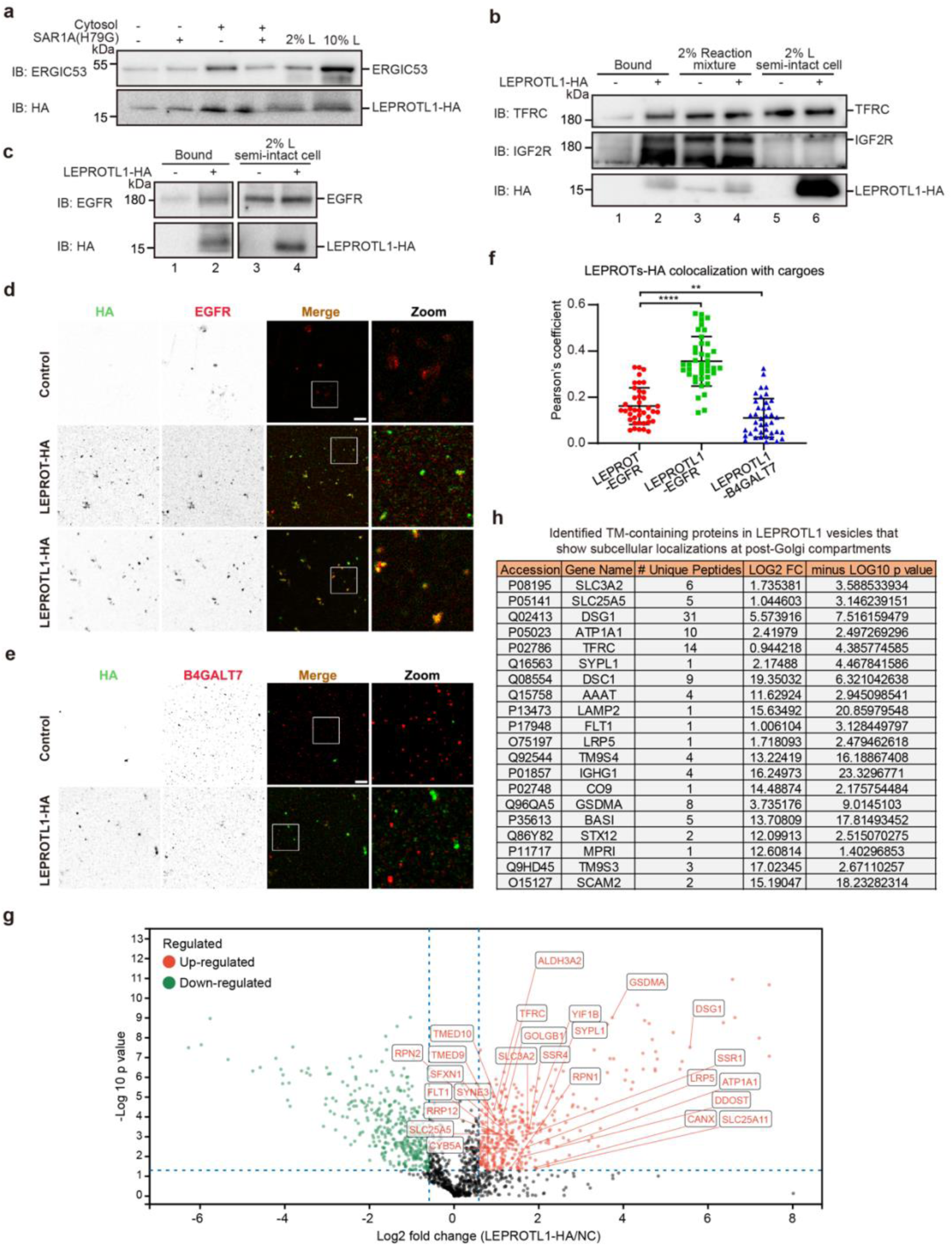
LEPROTs bud into COPI vesicles with specific clients. (**a**) Analyses of the effects of SAR1A(H79G) on the budding efficiency of the indicated cargo proteins. The vesicle formation assay was performed using HEK293T cells expressing LEPROTL1-HA in the presence of the indicated reagents, including SAR1A (H79G) or rat liver cytosol. Vesicle fractions were then analyzed by immunoblotting. (**b,c**) Analyses of the presence of EGFR, TFRC, and IGF2R in the immuno-isolated LEPROTL1-HA vesicles. The vesicle formation assay was performed using untransfected HEK293T cells or cells transfected with plasmids encoding LEPROTL1-HA. LEPROTL1-enriched vesicles were immunoprecipitated by anti-HA M2-agarose affinity beads and analyzed by immunoblotting. Notably, LEPROTL1-HA was mostly in the dimeric form in the immunoisolated vesicle fraction. The reaction mixture was a mixture of anti-HA beads with the medium speed supernatant following the vesicle formation assay. (**d,e**) Fluorescent analysis of the presence of EGFR (**d**) or B4GALT7 (**e**) in the LEPROT-HA or LEPROTL1-HA vesicles. The vesicle formation assay was performed using untransfected HEK293T cells or cells transfected with plasmids encoding LEPROT-HA or LEPROTL1-HA. LEPROT- or LEPROTL1-enriched vesicles were incubated with coverslips pre-incubated with poly-L-lysine and analyzed by immunofluorescence. LEPROT-HA and LEPROTL1-HA were stained with HA antibody (green), EGFR was stained with EGFR antibody (red), and B4GALT7 was stained with B4GALT7 antibody (red). Scale bar = 1 μm. (**f**) Quantification of the Pearson’s correlation coefficients for LEPROTs and the EGFR or B4GALT7 signal. Values are mean ± SD; n=40. ****p<0.0001, **p<0.01, two-tailed Student’s t-test. Scale bar = 1 μm. (**g**) Label-free quantitative mass spectrometric analysis to profile the vesicles enriched with LEPROTL1-HA. The vesicle formation assay was performed using untransfected HEK293T cells (the non-specific group, NC) or cells transfected with plasmids encoding LEPROTL1-HA. Vesicle fractions were then immuno-isolated by anti-HA agarose affinity beads. The proteins in the vesicle fractions were trypsin-digested and analyzed by label-free mass spectrometry. The average log2 fold changes of the identified proteins in the LEPROTL1-HA group compared to the NC group were calculated and plotted against the minus log10 P-value. TM-containing proteins identified in the LEPROTL1-HA vesicles are highlighted. (**h**) The list of identified transmembrane proteins located at the plasma membrane, endosomes, or lysosomes.

### LEPROTs interact with a subset of TM-containing proteins

Subsequent to our vesicle formation assays, we isolated vesicles enriched with LEPROT or LEPROTL1 using anti-HA conjugated beads. Untransfected HEK293T cells were used as a negative control. Immuno-blotting revealed that vesicles enriched with LEPROTL1 contained EGFR, TFRC (known as transferrin receptor protein 1 or TfR1), and IGF2R (**Fig. 3b** and **Fig. S3d**, compare lanes 1 and 2), whereas those with LEPROT contained TFRC and IGF2R, but not EGFR (**Fig. 3c** and **Fig. S3e**, compare lanes 1 and 2). In further experiments, we immune-stained the isolated vesicles and measured the fluorescence to analyze the co-packaging of LEPROTs with specific TM-containing proteins. Many of the EGFR signals overlapped with LEPROTL1 with significantly more colocalization than with LEPROT (**Fig. 3d,f**). When B4GALT7, a TM-containing Golgi resident protein, was tested, no detectable overlap was seen with LEPROTL1 (**Fig. 3e,f**). These results confirm that LEPROTs package some TM receptors into COPI vesicles.

To uncover additional LEPROTs clients in these vesicles, we subjected the immuno-isolated vesicles to label-free quantitative mass spectrometry. Peptides from three biological repeats led to the identification of 26 and 51 TM-containing proteins in LEPROT and LEPROTL1-enriched vesicles, respectively, with an enrichment factor greater than 1.7-fold (**Fig. 3g, Fig. S3f**, **Table S2** and **S3** sheets 3 and 5). Notably, the majority of these proteins, such as TFRC, were verified by the Uniprot database as being associated with post-Golgi subcellular localization, including the PM, endosome, and lysosomes (**Fig. 3h** and **Fig. S3g**). In addition, a subset of TM-containing clients were ER-resident proteins that were likely subjected to COPI-dependent retrieval. We also found proteins that were identified with high confidence in the LEPROT-enriched groups but not in the corresponding control groups (**Table S2** and **S3** sheet 4). These could be potential clients for LEPROTs pending further verification.

Notably, a substantial number of the identified proteins in these proteomics experiments are not TM-containing proteins. Most of these proteins are cytosolic factors potentially involved in regulating LEPROTs-mediated vesicle formation. For soluble cargo within the endomembrane system, we speculate that LEPROTs may regulate the COPI packaging of some TM-containing co-adaptors or co-receptors, which in turn capture luminal contents for COPI vesicles.

When comparing the client selectivity of the two LEPROTs, we noticed that some of the identified proteins were common to both vesicle types, whereas others were unique to each (**Fig. 3h** and **Fig. S3g**). The fact that LEPROTL1 engaged more hits than LEPROT implies a more active role in regulating COPI trafficking, which is in line with the observation that LEPROTL1 deletion results in a stronger Golgi phenotype than LEPROTs deletion (**Fig. 1e-g**). Taken together, these results indicate that LEPROTs-enriched vesicles co-package with specific clients destined for post-Golgi compartments, highlighting distinct cargo profiles for each protein variant. Given that LEPROTs have minimal cytosolic or luminal domains, we anticipated that LEPROTs engage their clients via TM domains. First, we tested whether LEPROTs are associated with known Golgi-resident or -passing proteins. As expected, endogenous IGF2R, EGFR, or TFRC, which are all TM domain-containing receptors, were clearly found in anti-HA precipitates when using cells expressing LEPROTL1-HA (**Fig. S2d**). In contrast, none of the known soluble proteins inside or outside the Golgi, including cathepsin D, SHH, and GAPDH, or B4GALT1, an irrelevant Golgi membrane protein, were associated with LEPROTL1 (**Fig. S2d**). Notably, the levels of potential clients for LEPROTs are not altered upon LEPROTs deletion (**Fig. S2e** and **Table S1**).

Next, we tested interactions between TM receptors and COPI coats, which are possibly bridged by LEPROTs. As expected, purified GST-tagged COPD, when incubated with cell lysates and precipitated, was able to pull down sufficient amounts of EGFR (**Fig. 4a,b**), confirming some capture of passing TM receptors by COPI vesicles. Consistently, deletion of LEPROTs in cells largely diminished such interactions (**Fig. 4a,b**). The residual COPD-EGFR binding in the absence of LEPROTs implies the existence of additional mechanisms for EGFR detainment by COPI coats. As a control, Golgi-resident integral membrane protein GOLIM4, a known client of GOLPH3 (13), also bound to COPD, but the association was not LEPROT-dependent (**Fig. 4a,c**). These results suggest that LEPROTs serve as cargo adaptors for TM receptors.

**Fig. 4.**
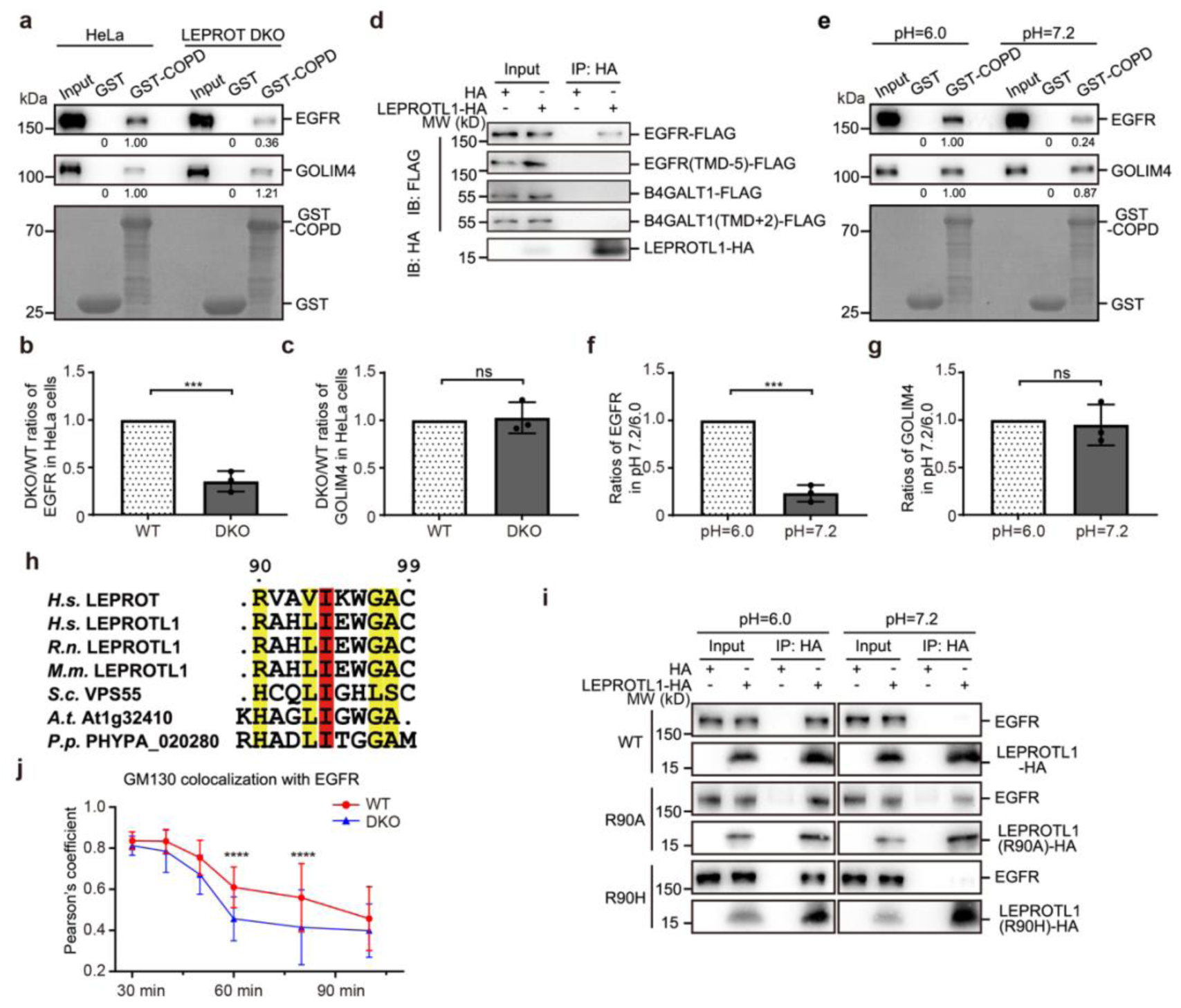
LEPROTs are COPI adaptors for TM receptors. (**a**) Western blot showing GST pull-down with endogenous EGFR and GOLIM4 in wild-type or LEPROT DKO HeLa cells. Quantifications of EGFR, GOLIM4 levels (normalized by their respective input protein levels) are also shown. (**b,c**) Quantitative data for (**a**) as mean ± SD (n=3). ***p<0.001, ns: not significant, two-tailed Student’s t-test. (**d**) Co-immunoprecipitation experiments showing the interaction between EGFR-FLAG, EGFR (TMD-5)-FLAG (LGIGL), B4GALT1-FLAG, B4GALT1 (TMD+2)-FLAG (AV) and LEPROTL1 in pH 6.0 MES buffer containing 1% digitonin. (**e**) Western blot showing GST pull-down with endogenous EGFR and GOLIM4 in pH 6.0 or 7.2 MES buffer. Quantifications of EGFR, GOLIM4 levels (normalized by their respective input protein levels) are also shown. (**f,g**) Quantitative data for (e) as mean ± SD (n=3). ***p<0.001, ns: not significant, two-tailed Student’s t-test. (**h**) Multispecies alignment of LEPROT 3-4 loop between *Homo sapiens (H.s.), Rattus norvegicus (R.n.), Mus musculus (M.m.), Saccharomyces cerevisiae (S.c.), Arabidopsis thaliana (A.t.),* and *Physcomitrium patens (P.p.)*. (**i**) Co-immunoprecipitation experiments showing the interaction between LEPROTL1 wild-type or R90D, R90H mutant and EGFR in pH 6.0 or 7.2 MES buffer containing 1% digitonin. (**j**) Quantification of the Pearson’s correlation coefficients in Fig. S4a for GM130 and HA-EGFR signals at different biotin addition times. Values are mean ± SD; n=50. ****p<0.0001, two-tailed Student’s t-test. Scale bar = 10 μm.

To probe the TM selectivity imposed by LEPROTs proteins, we compared the TM length of LEPROTs with EGFR and B4GALT1. The four TM domains of LEPROTs each contain ∼21 a.a. on average, implying a potential preference for relatively longer TM domains. TM domains in membrane proteins that have post-Golgi localization have been proposed to be longer in general than proteins residing in Golgi (37, 38). EGFR-TM has 23 a.a. and B4GALT1-TM is 19 a.a. in length. As expected, truncating EGFR-TM to 19 residues abolished its interactions with LEPROTL1 (**Fig. 4d**) though the mutant retained PM localization (**Fig. S2f**), which ensures the possibility of meeting with LEPROTL1. However, extending B4GALT1-TM to 21 residues did not have this effect (**Fig. 4d**), and the mutant resided mostly in the Golgi, similar to the wild-type (**Fig. S2f**). These analyses reveal that TM length can be read by LEPROTs when selecting clients. The results also indicate that additional factors are involved in the process. All of the TM lengths in candidates identified in the vesicle formation assay are equivalent to or longer than 21 a.a., supporting the notion that LEPROTs choose clients based on, at least in part, the length of their TM domains.

We reason that LEPROTs aim to grab clients at medial to trans Golgi before they exit the apparatus and release them at cis Golgi upon COPI retrieval. Therefore, LEPROTs are expected to have higher affinity for clients at trans Golgi than cis Golgi. To this end, we tested whether COPD-EGFR interactions are pH-sensitive, because Golgi exhibit a gradual decrease in pH from cis to trans (39, 40). COPD pulled down more EGFR from HeLa cell lysates at pH 6.0 than at pH 7.2, and COPD-GOLIM4 interactions were not sensitive to pH (**Fig. 4e-g**). As expected, COPD-LEPROTL1 interactions, which occur in the cytosol, were not pH-dependent (**Fig. S2g,h**), and LEPROTs deletions exerted no detectable impact on the existing pH gradients in the ER and Golgi (**Fig. S2i,j**). Collectively, LEPROTs likely detain TM receptors at trans Golgi via COPI-coated vesicles and drop them off at cis Golgi.

LEPROTs have only a few residues in either the 1-2 loop or 3-4 loop that are accessible to the Golgi lumen. We identified a conserved Arg (His in lower eukaryotes) in the 3-4 loop of LEPROTL1 that potentially senses pH changes (**Fig. 4h**). To probe the role of R90, we replaced it with a H to boost the pH sensitivity or an A to disrupt it. We then analyzed interactions between LEPROTL1 mutants and EGFR at various pH. Similar to the wild-type, R90H was capable of engaging EGFR at pH 6.0, but not pH 7.2 (**Fig. 4i**). In contrast, R90A interacted with EGFR at both pH 6.0 and 7.2. These results suggest that R90 controls pH-dependent unloading of LEPROTs clients.

### LEPROTs regulate Golgi retention of TM receptors

To compare the intracellular pathways and dynamics of TM receptors in wild-type and DKO cells, we utilized a well-established “<u>r</u>etention <u>u</u>sing <u>s</u>elective <u>h</u>ooks” (RUSH) system (41). In this system, cargo proteins were initially trapped in the ER and released synchronously upon biotin addition. As a typical PM-enriched TM receptor, EGFR would exit the ER towards the Golgi and leave the Golgi for the PM. When RUSH-competent EGFR was released from the ER, it mostly reached the Golgi within 30 min. Subsequent PM delivery could be seen after 60 min. To measure Golgi retention using this cargo, we analyzed Pearson’s coefficient between EGFR signals and Golgi marker GM130 over 30-100 min. Notably, the Golgi departure of EGFR was accelerated in DKO cells (**Fig. 4j** and **Fig. S4a**), which is consistent with a failure in sufficient Golgi retention. In contrast, the Golgi departure of soluble cargo, including secreted protein sonic hedgehog (SHH) and lysosomal enzyme cathepsin Z (CTSZ) was not affected by the deletion of LEPROTs (**Fig. S4b-e**), ruling out soluble proteins being the clients of LEPROTs. These results indicate that LEPROTs are capable of retaining certain proteins in Golgi.

We also analyzed the intracellular localization of Golgi-resident proteins in the presence or absence of LEPROTs. IGF2R (also known as cation-independent mannose-6-phospate receptor [CI-M6PR]) is a receptor for both insulin-like growth factor 2 and lysosomal enzymes modified with mannose-6-phosphate, and it recycles between the Golgi and the cell surface (42). IGF2R was clearly found in Golgi in wild-type HeLa cells but largely dispersed into endosomes, instead of lysosomes, when LEPROTL1 or both LEPROTs were deleted (**Fig. S5a-c**). Consistent with a failure in Golgi retention, the intracellular signals of IGF2R covered larger areas and became distant from Golgi (**Fig. S5d,e**). As a result, cathepsin D that are sorted to lysosome by IGF2R suffered processing defects in these mutant cells (**Fig. S5f**). The same results were obtained in LEPROT DKO COS-7 cells (**Fig. S5g-i**). The localization defects of IGF2R could be restored when LEPROTL1 was expressed in DKO cells (**Fig. S5j-m**). By comparison, GALNT2, a TM-containing Golgi enzymes and known clients of GOLPH3, did not leave the Golgi in LEPROT DKO cells (**Fig. S5n**). Consistently, LEPROT deletion had no impact on COPII-mediated ER export for EGFR, SHH or CTSZ (**Fig. S4f-h**). The levels of major coats for various protein trafficking pathways was not significantly altered upon deletion of LEPROTs (**Fig. S2k** and **Table S1**). These results reinforce the notion that LEPROTs work with COPI for Golgi retention of TM receptors.

### LEPROTs regulate modification and activities of TM receptors

Next, we tested the levels of known Golgi modification, including palmitoylation and O-glycosylation, using EGFR and TFRC as representative receptors in the context of LEPROTs. We measured the levels of EGFR glycosylation, specifically O-Gal modification, which can be isolated by vicia villosa agglutinin (VVA) pull-down, and found a decrease in DKO HeLa cells (**Fig. 5a,b**). Similar results were obtained in U-2 OS cells, though the signals were much weaker due to a lower level of endogenous EGFR (**Fig. 5c,d**). These changes are less likely due to altered expression of O-Gal modification enzymes as GALNT2, one of the major O-Gal-modifying enzymes, was strongly elevated in response to LEPROTs deletion in HeLa cells, and remained unchanged in U-2 OS cells (**Fig. S6a,b and Table S1**). Importantly, the deletion of LEPROTs did not cause a global reduction in O-Gal modification (**Fig. S6c**). We also tested whether over-expression of LEPROTs causes the opposite effects. Consistently, O-Gal addition was promoted in LEPROTL1-transfected cells (**Fig. S6d,e**). When comparing wild-type or LEPROT DKO HeLa cells, the palmitoylation levels of TFRC were reduced as expected upon deletion of LEPROTs (**Fig. 5e,f**), consistent with insufficient Golgi retention of the receptor. The same results were obtained with U-2 OS or HepG2 cells (**Fig. S6f-i**) and for EGFR (**Fig. S6j,k**). Finally, we tested whether TM receptor processing is also altered in a previously identified COPI adaptor. As expected, O-Gal modification of EGFR did not result in detectable changes in GOLPH3 depletion (**Fig. S6l,m**), even when glycosylation enzymes had defects in Golgi retention(13), suggesting that the retention of clients themselves, but not the corresponding enzymes, is more important for proper modification of TM receptors. Collectively, these data demonstrate that LEPROTs specifically influence TM receptor processing by regulating their Golgi retention.

**Fig. 5.**
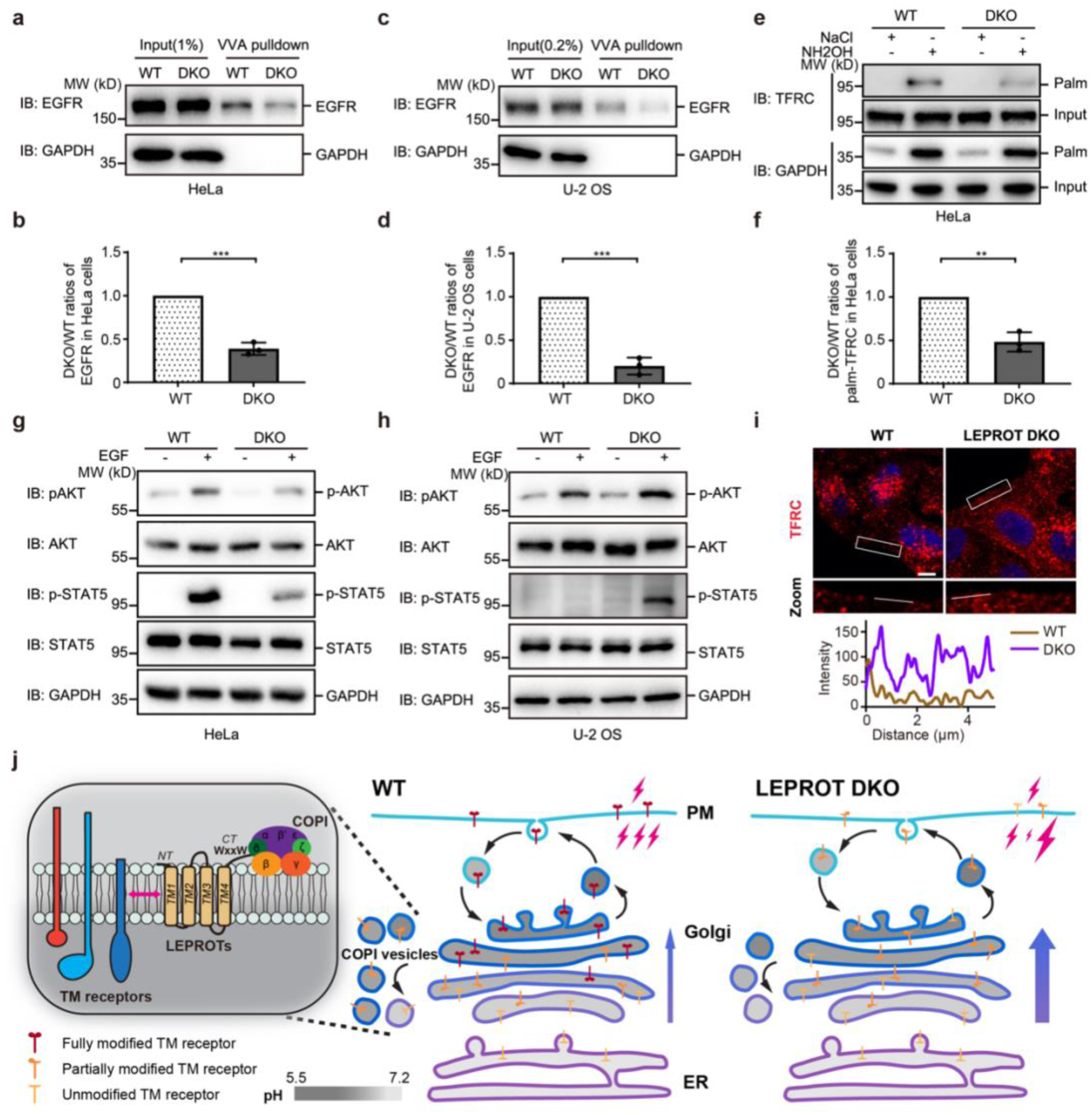
LEPROTs regulate TM receptor processing and activity. (**a,c**) Wild-type and LEPROT DKO HeLa or U-2 OS cell lysates were subjected to VVA pull-down assay. (**b,d**) The quantitative data of (**a,c**) shown as mean ± SD (n=3). ***p<0.001. (**e**) Wild-type and LEPROT DKO HeLa cells were subjected to acyl-RAC assay for palmitoylation detection. (**f**) The quantitative data of E shown as mean ± SD (n=3). **p<0.01. (**g, h**) Wild-type and LEPROT DKO HeLa or U-2 OS cells were treated with 10 ng/ml EGF for 10 min and harvested for Western blot using the indicated antibodies. (**i**) Wild-type and LEPROT DKO HeLa cells were stained with TFRC antibody. The plasma membrane areas are in the zoomed-in images. The white line indicates the pixels used for the fluorescence intensity profile plots. Nuclei were indicated using DAPI (blue). Scale bar = 10 μm. (**j**) A model of the role of LEPROTs in the sufficient modification of TM receptors.

Golgi processing of TM receptors greatly impacts their activity and function. O-Gal modification of EGFR has been suggested to be tightly associated with its oncogenic activity (43, 44). Thus, we treated wild-type or DKO cells with EGF and monitored the signaling strength. Phosphorylation of AKT and STAT5 are considered downstream events of EGFR signaling (45, 46). We found that, in HeLa cells, deletion of LEPROTs caused a weakened EGFR signal (**Fig. 5g**). In contrast, U-2 OS DKO cells exhibited an elevated EGF response compared to wild-type cells (**Fig. 5h)**. To confirm that the altered EGFR signal is largely linked to levels of O-Gal, we transfected DKO cells with GALNT2. Overexpression of GALNT2 in U-2 OS DKO cells successfully repressed EGFR activation as expected (**Fig. S6n**). Palmitoylation of TFRC has been shown to play a role in receptor endocytosis (47). When TFRC localization was compared in wild-type and DKO cells, the presence was increased in the PM in DKO cells (**Fig. 5i**), accompanied by a decreased overlap with endosomes (**Fig. S6o,p**) and coinciding with reduced palmitoylation of TFRC in these cells. Taken together, the findings indicate that LEPROTs activity is associated with TM receptor activity by controlling their Golgi processing, the physiological outcome of which can be cell-type specific.

## DISCUSSION

The LEPROT gene was initially identified as an alternatively spliced transcript of the leptin receptor gene (48). Subsequently, the homologous gene for LEPROTL1 was cloned (49). However, the function of the LEPROT family of proteins is poorly understood. Our results demonstrate that LEPROTs serve as a COPI adaptor by bridging Golgi-passing TM receptors and COPI coats (**Fig. 5j**). Thus, LEPROTs and leptin receptor are superficially irrelevant, but their paths incidentally reconnect in the Golgi.

The LEPROT protein family is conserved throughout eukaryotic cells. In yeast, the homologous protein VPS55 was thought to mediate late endosome to vacuole trafficking (50). Consistently, VPS55 localizes mostly to endosomes but not Golgi (50, 51). Sequence comparisons of high eukaryote LEPROTs and VPS55 suggest that the four tandem TMs are largely conserved, but the C-termini of these proteins diverge. We identified a highly conserved WxxW motif in the C-termini of vertebrate LEPROTs. The motif is critical for the direct interaction of LEPROTs with COPI coats and, thus, a new localization preference and a new role for the proteins. Such a motif is missing in VPS55, which could explain its endosomal, but not Golgi, enrichment. The differentiated roles of the LEPROT family members implicate different demands and dynamics of protein trafficking in different cells. Nevertheless, it is yet to be determined whether the mammalian LEPROTs possess some function in regulating endosomes to Golgi trafficking.

The LEPROT class of COPI adaptors differs in several aspects from known adaptors, including Vps74p and GOLPH3. Vps74p (or GOLPH3) is a cytosolic, soluble adaptor bridging the coatmer and cytosolic tail of Golgi-localized enzymes (22). In contrast, the LEPROTs are membrane-embedded, linking COPI coats to TM receptors; the interactions with COPD are solvent-exposed but the recruitment of clients is most likely TM-based. The actions of Vps74p and GOLPH3 are largely expected, as Golgi-localized glycosylation enzymes are supposed to remain in the Golgi. On the other hand, the role of LEPROT, which delays Golgi exit of TM receptors, is uncertain until proven. We showed that COPD interacts with EGFR in a LEPROT-dependent manner, confirming the existence of some COPI-mediated retention for receptors that are transiting through the Golgi. Finally, the mechanism for client loading and release by LEPROTs coincides with the pH gradients through Golgi stacks. This workflow is not found in Vps74p/GOLPH3-mediated COPI trafficking, in which PI4P-binding is involved. Notably, in the absence of both LEPROTs, EGFR retained a weak association with COPD, suggesting the existence of additional COPI adaptors that possess similar activity as LEPROTs.

In addition to TM receptors, LEPROTs may facilitate retrieval of escaped ER-resident membrane proteins in the Golgi. The comparison of identified clients for LEPROTs revealed that these adaptors recognize the length of TM domains and preferentially engage relatively longer TMs. LEPROTs may also indirectly affect luminal cargo via co-adaptors that are yet to be identified. The two LEPROTs appear to be functionally redundant when expressed at a sufficient level, but each may be in charge of a unique set of clients. Alternatively, endogenous LEPROT may have more endosomal localization than LEPROTL1 and, thus, exerts additional functions there as reported previously (29). Overall, our deletion analysis suggests that LEPROTL1 is more active than LEPROT in serving as a COPI adaptor, at least in the setting of HeLa cells.

We showed that mammalian LEPROTs delay the Golgi departure of TM receptors, ensuring their sufficient processing, including O-Gal addition and palmitoylation. Notably, transgenic mice expressing human LEPROTs exhibited limited PM exposure of growth hormone receptor (GHR) and subsequent decrease in growth hormone action, consistent with boosted Golgi retention of the receptor (52). LEPROTs have also been implicated in tumorigenesis, many of which can now be linked to regulation of growth factor receptor (53–58). Though the sufficient Golgi retention and accompanied processing of TM receptor is ensured in the presence of LEPROTs, the consequences of such an action are likely cellular context-dependent. Notably, the LEPROT-mediated retention of clients is weak and temporary in nature. It is not an absolute blockade of Golgi exit for these clients, but has an evident impact on their processing. Our results provide important insights into the physiological role of COPI trafficking and potential therapeutic strategy that targets manipulation of TM receptors.

## METHODS

### Plasmids

Full-length cDNA of human LEPROT, LEPROTL1, COPD, GALNT2, SHH, B4GALT1, CTSZ and EGFR were cloned from a cDNA library prepared from HEK293T or U-2 OS cells. LEPROTL1 truncations and mutations were generated using overlap PCR. These PCR products were cloned into pcDNA4/TO vector (Invitrogen), pGEX-6p vector (GE Healthcare), or pLenti-GFP-Puro-CMV vector (Addgene, 17448) with the indicated tags. sgRNAs were cloned into px458 vector (Addgene, 48138).

### Cell culture, transfection, and immunofluorescence

HeLa cells, COS-7 cells, U-2 OS cells, and HepG2 cells were grown in complete DMEM with 10% FBS at 37 °C in 5% CO_2_. Plasmids were transfected with Lipofectamine 3000 Transfection Reagent (Invitrogen) according to the manufacturer’s protocol. Cells grown on coverslips were fixed with 4% polyoxymethylene for 15 min at room temperature, permeabilized with 0.1% Triton X-100 in PBS for 10 min at room temperature, and then blocked with 3% BSA in PBS for 1 h at room temperature. Primary antibodies were diluted in 3% BSA in PBS and incubated overnight at 4 °C or at room temperature for 1 h. Secondary antibodies conjugated with Alexa Fluor 488, 568, or 647 (1:500) were diluted in 3% BSA in PBS and incubated for 1 h in the dark at room temperature. Images were taken on a Nikon A1R+ confocal microscope. The following primary antibodies were used: anti-B4GALT1 (Abcam, ab121326), anti-B4GALT7 (Proteintech, 10535-1-AP); anti-EEA1 (BD Biosciences, 610457), anti-GM130 (BD Biosciences, 610823), anti-HA (Roche, 11867423001), anti-Golgin-97 (Abcam, ab84340), anti-IGF2R (Abcam, ab124767), anti-IGF2R (Abcam, ab2733), anti-PDI (Abcam, ab2792), anti-LAMP1 (Cell Signaling Technology, 9091S), anti-LAMP1 (Abcam, ab25630), anti-TFRC (Abcam, ab269513), anti-FLAG (Sigma, F3165), anti-SEC31A (BD Biosciences, 612351), anti-calreticulin (Proteintech, 27298-1-AP), anti-GM130 (Proteintech, 11308-1-AP) and anti-COPB1 (Invitrogen, PA1-061). Various Alexa Fluor-conjugated secondary antibodies were purchased from Invitrogen. GLIM analysis was performed as described previously(32).

### Cell treatments

HeLa/HepG2/U-2 OS or siRNA transfected cells were treated with 10 ng/ml EGF for 10 min and analyzed by SDS-PAGE and immunoblotting to determine the phosphorylation level of key proteins of the corresponding signaling pathways.

### Western blot

Proteins were extracted with lysis buffer (25 mM Tris-HCl, pH 7.4, 150 mM NaCl, 1% NP-40, 1 mM EDTA, 5% glycerol) containing protease and phosphatase inhibitors and analyzed by SDS-PAGE and Western blot. Proteins were detected using the following specific antibodies: anti-HA (Huaxingbio, HX1820), anti-ERGIC-53 (Proteintech, 13364-1-AP), anti-β-tubulin (Santa Cruz, sc-101527), anti-α-tubulin (Proteintech, 66031-1-Ig), anti-BiP (BD Biosciences, 610979), anti-GOLIM4 (Abcam, ab197595), anti-cathepsin D (CTSD; Proteintech, 21327-1-AP), anti-COPA (Santa Cruz, sc-398099), anti-COPB1 (Invitrogen, PA1-061), anti-COPB2 (Abclonal, A21294), anti-COPD (Abcam, ab96725), anti-COPE (Proteintech, 11457-1-AP), anti-COPG1 (Proteintech, 12393-1-AP), anti-COPZ1 (Abclonal, A12148), anti-HA (Cell Signaling Technology, 3724S), anti-SHH (Cell Signaling Technology, 2207S), anti-GAPDH (BBI Life Sciences, D110016-0200), anti-EGFR (PTM Biolabs, PTM-5438), anti-AKT (Cell Signaling Technology, 9272S), anti-pAKT (Ser473) (Cell Signaling Technology, 4051S), anti-STAT5 (Cell Signaling Technology, 94205T), anti-pSTAT5 (Tyr694) (Cell Signaling Technology, 4322T), anti-REEP5 (Proteintech, 14643-1-AP), anti-SEC31A (BD Biosciences, 612351), anti-β-actin (Proteintech, 66009-1-Ig), anti-GFP (Roche, 11814460001), anti-TFRC (Santa Cruz, sc-65882), anti-AP1G1 (BD Biosciences, 610385), anti-ARF1 (Proteintech, 10790-1-AP), anti-GOLPH3+GOLPH3L (Proteintech, 19112-1-AP), anti-IGF2R (Abcam, ab124767), anti-GALNT2 (Novus Biologicals, NBP1-83394), and anti-LEPROTL1 (Sangon Biotech, immunizing rabbits with synthetic peptides corresponding to the C-terminus of human LEPROTL1). Blots were visualized using HRP substrate and captured by an imaging system (4600SF, Tanon).

### Genome editing using CRISPR/Cas9

LEPROT KO, LEPROTL1 KO, LEPROT DKO HeLa/COS-7/HepG2/U-2 OS, GOLPH3 KO HeLa cells, and LEPROT-2xHA knockin HeLa cells were constructed using the CRISPR/Cas9 system. CRISPR-targeting oligonucleotides designed for LEPROT, LEPROTL1, or GOLPH3 were inserted into the pX458 vector. Cells were transfected with corresponding pX458 vectors, and GFP-positive cells were sorted as single clone into a 96-well plate using a BD Influx cell sorter (BD BioSciences). Knockout and knockin cells were identified by sequencing. Genomic DNA was extracted using DNA extraction buffer (10 mM Tris-HCl, pH 8.2, 50 mM KCl, 2.5 mM MgCl2, 0.45% NP-40, 0.45% Tween-20 and 0.01% gelatin, proteinase K). Gene-specific PCR products were amplified by M5 HiPer plus Taq (Mei5 Biotech, MF002plus) and cloned into pCI-neo (Promega) vector for sequencing. The detailed information on sgRNA sequences, protein changes, and cDNA changes in KO cell lines is listed in **Table S4**.

### Alkaline extraction assay

Briefly, LEPROTs-2xHA (KI) HeLa cells were washed with pre-chilled PBS, subsequently suspended in Buffer A (Tricine 20 mM, sucrose 250 mM, pH 7.8, and protease inhibitor cocktails), and kept on ice for 20 min. The cells were homogenized before centrifugation at 3,000 *g* for 10 min to remove the nucleus. The supernatant was considered to be the total lysate. The total lysates were further centrifuged at 100,000 *g* for 40 min at 4 °C to obtain the cytosol and total membranes of all organelles. The total membrane fraction was subjected to alkaline extraction (0.1 M Na_2_CO3, pH 11) followed by centrifugation at 100,000 *g* for 40 min. Both the pellets and supernatant were analyzed by SDS-PAGE and immunoblotting.

### MP modification assay

HeLa cells transfected with LEPROTL1 constructs were washed in PBS and then treated with 0.05% digitonin for 10 min at 4 ℃. Samples were incubated for 30 min at 4 ℃ with 1 mM maleimide PEG 5 kDa (MP) in the absence or presence of 20 mM DTT. All samples were treated with 20 mM DTT and 1% Triton X-100 before analysis by SDS-PAGE and immunoblotting.

### Co-immunoprecipitation

For COPI immunoprecipitation, cells were lysed in 1% NP-40 IP buffer (25 mM Tris-HCl, pH 7.4, 150 mM NaCl, 1 mM EDTA, 5% glycerol, and protease inhibitor cocktails). For membrane receptor protein immunoprecipitation, cells were lysed in 1% digitonin IP buffer (25 mM Tris-HCl, pH 7.4, 150 mM NaCl, 1 mM EDTA, 5% glycerol, and protease inhibitor cocktails). Cell lysates were incubated with anti-HA agarose (Sigma, A2095) for 2 h at 4 ℃. Washed precipitates were separated by SDS-PAGE and immunoblotting.

### Protein purification

GST-tagged human COPD proteins were expressed in *Escherichia coli* BL21 (DE3). Harvested cell pellets were lysed by ultrasonication in protein purification buffer (25 mM Tris-HCl, pH 7.4, 300 mM NaCl, and protease inhibitor cocktails). Bacterial lysates were centrifuged at 30,000 *g* for 1 h at 4 ℃. The supernatant was incubated with pre-washed GST-Sefinose Resin (BBI Life Sciences, C600031) for 1 h at 4 ℃. Resins were washed with washing buffer A (25 mM Tris-HCl, pH 7.4, 300 mM NaCl), and then the target protein was eluted with 10 mM glutathione. Next, the GST fusion tag were removed by the 3C protease.

### Peptide pull-down assay

Biotinylated LEPROTL1 2-3 loop or CT peptides were synthesized by Sangon Biotech and dissolved in H2O or 200 mM Tris-HCl (pH 8.0). For each sample, 20 µM peptide was incubated with 25 µl of pre-washed streptavidin resin (Thermo Fisher Scientific) for 1 h at 4 ℃. Beads bound with peptides were mixed with 500 µg of HeLa cell lysate in lysis buffer (25 mM Tris-HCl, pH 7.4, 150 mM NaCl, 1% NP-40, 1 mM EDTA, 5% glycerol) containing protease and phosphatase inhibitors or 1 µg of purified COPD and incubated for 2 h at 4 ℃. Next, the samples were washed extensively with lysis buffer and analyzed by SDS-PAGE and immunoblotting.

### RUSH assay

HeLa or LEPROT DKO HeLa cells were grown on 15-mm glass coverslips in 24-well culture dishes for 24 h. Then cells were transfected with plasmids encoding Str-KDEL and SBP-HA-EGFR, SBP-HA-SHH, or SBP-GFP-CTSZ using Lipofectamine 3000 Transfection Reagent (Invitrogen) according to the manufacturer’s protocol. After 24h transfection, RUSH assays were performed by treating cells with complete medium containing 40 μM biotin (Sigma-Aldrich) and 100 ng/μl cycloheximide (Sigma-Aldrich) for the indicated time. In parallel, a control sample without biotin was used to monitor the leakiness of the RUSH construct-encoded cargo protein before the release (0 min). After the release, cells were fixed at the indicated time points. Images were taken by a Nikon A1R+ confocal microscope.

### Vesicle formation assay

In vitro vesicular release assays were performed using wild-type and LEPROT DKO HeLa cells. The approach was a modified version of the previous vesicle budding reaction(10, 59). HeLa cells grown in one 10-cm dish until approximately 90% confluence were permeabilized in 5 ml of ice-cold KOAc buffer (110 mM potassium acetate, 20 mM HEPES, pH 7.2, 2 mM magnesium acetate) containing 40 mg/ml digitonin on ice for 5 min. The semi-intact cells were then sedimented by centrifugation at 300 *g* for 3 min at 4 ℃. The cell pellets were resuspended in 5 ml of high salt KOAc buffer (0.5 M potassium acetate) and incubated on ice for 5 min. After incubation, the permeabilized cells were washed twice with 1 ml of KOAc buffer and resuspended in 200 ml of KOAc buffer. The budding assay was performed by incubating semi-intact cells (∼0.02 OD/reaction) with 2 mg/ml of rat liver cytosol in a 100 ml reaction mixture containing 200 mM GTP and an ATP regeneration system (40 mM creatine phosphate, 0.2 mg/ml creatine phosphokinase, and 1 mM ATP). After incubation at 32 ℃ for 1 h, the reaction mixture was centrifuged at 14,000 *g* to remove cell debris and large membranes. The medium-speed supernatant was then mixed with 60% OptiPrep to 35% OptiPrep and overlaid with 1 ml of 30% OptiPrep and 50 ml of KOAc buffer. The samples were centrifuged at 54,000 rpm in a S55S rotor in a Himac ultracentrifuge at 4 ℃ for 1.5 h. The 200 μl top fraction (i.e., vesicle fraction) was then centrifuged at 55,000 rpm to sediment small vesicles.

To immuno-isolate vesicles enriched with LEPROT and LEPROTL1, the volume of the reaction mixture was scaled up to 1 ml. The medium-speed supernatant was incubated with 10 ml of anti-HA agarose affinity beads at 4 ℃ overnight. After incubation, the beads were washed four times with 1 ml of KOAc buffer. The beads were then analyzed by immunoblotting or mass spectrometry.

To visualize the isolated vesicles, the vesicle fraction was diluted 1:1 with ice-cold KOAc buffer. The coverslips were incubated with 500 μl poly-L-lysine at 37 °C for 24 h. The diluted vesicle fraction was then incubated with the coverslips at 4 °C overnight and immunofluorescence performed as described. The vesicles were visualized by confocal microscopy and the images analyzed by Fiji.

### Preparing cytosol from HEK293T cells

HEK293T cells grown in two 15-cm dishes to ∼90% confluence were incubated in 500 μl ice-cold Buffer E (250 mM sorbitol, 50 mM Hepes, pH 7.2, 70 mM KOAc, 0.5 mM magnesium acetate, 5 mM EGTA-K) on ice and passed needle for 10 min. The collected lysates were centrifuged twice at 14,000 rpm for 10 min at 4 °C. The supernatant was then centrifuged at 55,000 rpm for 1 h at 4 °C. The top supernatant was collected as the “cytosol”. To deplete the COPI coat from the cytosol, Strep-Tactin Sepharose beads were incubated with the CT peptides for 1 h at 4 °C, and the cytosol was incubated with unprocessed and the CT peptide-incubated Strep-Tactin Sepharose beads for 2 h at 4 °C. The budding assay was performed using 2 mg/ml of cytosol from HEK293T cells.

### Sample preparation for mass spectrometry

The sample was analyzed by SDS-PAGE and the gel bands stained by Coomassie blue. The gel bands were cut into fragments and washed with 25 mM NH_4_HCO_3_/50% ACN. The gel fragments were shrank in ACN, swelled in 0.1 M NH_4_HCO_3_/ 10 mM DTT, and incubated at 55 °C for 45 min. Subsequently, 0.1 M NH_4_HCO_3_/ 55 mM ACN was added and the mixture incubated at room temperature for 45 min. The gel fragments were washed in 100 μl of 0.1 M NH_4_HCO_3_at room temperature for 15 min. The solution was removed and the gel fragments shrank in ACN to completely dry. The gel fragments were then swelled in 50 mM NH_4_HCO_3_/ 20 ng/μl trypsin on ice for 45 min. The solution was removed and replaced with 10 μl of 50 mM NH_4_HCO_3_, followed by incubation at 37 °C overnight. The tubes were centrifuged, the supernatants collected, and the peptides extracted with 25 mM NH_4_HCO_3_ at room temperature for 15 min and 5% formic acid/60% ACN twice. The extracts were combined and dried in a vacuum centrifuge. The sample was then reconstituted in 0.1% TFA for desalting using the Pierce C18 spin column. The columns were activated with 50% ACN in 0.1% TFA, and then equilibrated by 0.1% TFA. The sample solution was added on top of the resin bed and washed with 0.1% TFA equilibration solution. Samples were collected using elution buffer with 50% and 75% ACN in 0.1% TFA. Finally, the samples were dried in a Speedvac concentrator and stored at −20 °C for further mass spectrometric analysis.

### Mass spectrometric data analysis

The analyses presented in the volcano plots in Fig. 3g and Fig. S3f focused on proteins with abundances identified in all three biological replicates in both the NC group and the LEPROT or LEPROTL1 groups (Table S2, sheet 2 and Table S3, sheet 2). The abundance was normalized to the median value of each group. The average abundance of each identified protein was calculated from the normalized abundance values across the three replicates. A Student’s t-test was utilized to assess significant differences between the two experimental groups considering only changes for which the p-values were < 0.05 as significant.

We also analyzed proteins that were identified in all three replicates of the LEPROT or LEPROTL1 group but were absent in at least one replicate of the NC group (Table S2, sheet 4 and Table S3, sheet 4). For these proteins, missing abundance values were assigned a random number between 1 and 2000. After normalizing the data to the median value of each group, the average abundance of each protein was again calculated from the three replicates. Significant changes between the two experimental groups were determined using a Student’s t-test based on the abundance values from the three independent biological replicates, with p < 0.05 considered significant.

### Ratiometric pH measurement

Ratiometric pH measurements were performed as described previously (60). Briefly, experiments were performed on a Nikon A1R+ confocal microscope at 37 °C in 5% CO2. RpHLuorin2 was excited sequentially at 405 and 488 nm, and images were acquired with an emission wavelength at 532 nm.

### GST pull-down assay

The purified GST or GST tagged COPD protein (10µg) was incubated with 50 µl of prewashed GST-Sefinose Resin and incubated for 1 h at 4 ℃. Beads were washed with cold washing buffer B (50 mM MES, 150 mM NaCl, 1% digitonin, pH=6.0 or 7.2). 10-cm dish of confluent HeLa or LEPROT DKO HeLa lysates were added to the beads and allowed to incubate at 4 °C overnight. Then, the samples were washed extensively with washing buffer B and analyzed by SDS-PAGE and immunoblotting.

### VVA pull-down assay

HeLa lysates were collected using lysis buffer (25 mM Tris-HCl, pH 7.4, 150 mM NaCl, 1% NP-40, 1 mM EDTA, 5% glycerol) containing protease and phosphatase inhibitors. WT or DKO lysates were incubated with pre-washed Vicia Villosa Lectin Agarose beads (Vector Labs, AL-1233-2) with constant rotation at 4 ℃ overnight. Beads were washed with lysis buffer four times at 4 ℃ and analyzed by SDS-PAGE and immunoblotting.

### Palmitoylation detection assay

Palmitoylated TFRC was analyzed by acyl-resin-assisted capture (acyl-RAC) assay. Briefly, harvested cells were dissolved by ultrasonication in buffer B (100 mM HEPES, pH7.5, 1 mM EDTA, 2.5% SDS). Free thiols on unmodified cysteine residues on TFRC were blocked using 0.1% s-methyl methanethiosulfate (MMTS) at 42 ℃ for 15 min. Proteins were precipitated by pre-chilled acetone overnight at - 20 ℃, then washed three times with 70% cold acetone and resuspended in Buffer D (100 mM HEPES, pH 7.5, 1 mM EDTA, 1% SDS). Samples were then divided into two tubes and incubated with 0.24 M NH_2_OH or NaCl at room temperature for 20 min. Samples were incubated with pre-washed High Capacity Thiol Reactive Resin (Nanocs, AR-SS-2) with constant rotation at room temperature for 3 h. Resins were washed with Buffer D containing 8 M urea five times at room temperature and eluted with Buffer D containing 50 mM DTT at room temperature for 30 min. Eluted fractions were analyzed by SDS-PAGE and immunoblotting.

### Cell treatments

HeLa/HepG2/U-2 OS or siRNA transfected cells were treated with 10 ng/ml EGF or 500 ng/ml growth hormone for 10 min and analyzed by SDS-PAGE and immunoblotting to determine the phosphorylation level of key proteins of the corresponding signaling pathways.

### Transient siRNA-mediated knockdown

Lipofectamine RNAiMAX (Invitrogen, 13778150) was applied to transfect siRNA according to the manufacturer’s protocol. The siRNA sequences used were as follows: COPB1, 5’-AACUUCCUGGACUUCUGAUGA-3’;

### Statistical analysis

Images were measured using Fiji (Image J) or Imaris software (Bitplane, Switzerland). An image J plugin, JACoP (Just Another Co-localization Plugin), was used to evaluate Pearson’s correlation coefficients for two different signals. The data are presented as the mean ± SEM. Significance was determined by an unpaired two-sided Student’s t-test. Statistical analysis was graphically presented by GraphPad Prism 9 software.

### Data availability

All data generated or analyzed during this study are included in this article and its supplementary information files. Source data are provided with this paper. Proteomic datasets are deposited as accession codes.

## ACKNOWLEDGEMENTS

We thank Dr. Alicia Prater for proofreading, Drs. Sha Sun, Birong Shen for critical discussion, and Junying Jia, Yihui Xu, Jifeng Wang, Yun Feng, Chunliu Liu, Xixia Li, Zhongshuang Lv for technical support (Institute of Biophysics, CAS). JH is supported by grants from the National Natural Science Foundation of China (92254305 and 32230024), the Strategic Priority Research Program (XDB39000000), and Project for Young Scientists in Basic Research (YSBR-075) of the Chinese Academy of Sciences. JG is supported by grant from the National Natural Science Foundation of China (32100550).

## AUTHOR CONTRIBUTIONS

JH designed and supervised the project. JG performed most of the experiments with the help from YX, YW, and BY. CC and YG performed and analyzed the budding assay with mass spectrometry support from QJ and ZY. JH and YG wrote the manuscript with input from all authors.

